# Auditory brainstem responses are resistant to pharmacological modulation in Sprague Dawley wildtype and Neurexin1α knockout rats

**DOI:** 10.1101/2023.05.22.541775

**Authors:** Samuel Marashli, Philipp Janz, Roger Redondo

## Abstract

Sensory processing in the auditory brainstem can be studied with auditory brainstem responses (ABRs) across species. Although ABRs have been widely utilized to evaluate abnormalities in auditory brainstem physiology, there is limited understanding if ABRs can be useful tool to assess the effect of pharmacological interventions. Therefore, we set out to understand how pharmacological agents that target key transmitter systems of the auditory brainstem circuitry affect ABR physiology in rats. Given previous studies, demonstrating that Nrxn1α KO Sprague Dawley rats show substantial auditory processing deficits and altered sensitivity to GABAergic modulators, we used both Nrxn1α KO and wildtype littermates in our study. First, we probed how different commonly used anesthetics (isoflurane, ketamine/xylazine, medetomidine) affect ABRs waveforms. In the next step, we assessed the effects of different pharmacological compounds (diazepam, gaboxadol, retigabine, nicotine, baclofen and bitopertin) either under isoflurane or medetomidine anesthesia. We found that under our experimental conditions, ABRs are largely unaffected by diverse pharmacological modulation. Significant modulation was observed with i.) nicotine, affecting the late ABR components at 90 dB stimulus intensity under isoflurane anesthesia in both genotypes, and ii.) retigabine, showing a slight decrease in late ABRs deflections at 80 dB stimulus intensity, mainly in isoflurane-anesthetized Nrxn1α KO rats. Our study suggest that ABRs in anesthetized rats are resistant to a wide range of pharmacological modulators, which has important implications for the applicability of ABRs to study auditory brainstem physiology.

## 1 Introduction

Auditory brainstem responses (ABR), also known as brainstem auditory evoked potentials, are electrical potentials evoked by brief tones, which can be measured non-invasively and that speak to synaptic transmission within the auditory brainstem circuits. ABRs are widely used for assessing hearing thresholds^1^, intraoperative neuromonitoring^2^, screening for sensory abnormalities in neurodevelopmental disorders^3^, or testing ototoxicity in drug development^4^.

In both humans and rodents, ABRs consists of distinct deflections (also referred to as ‘waves’), that are generated by the activation of specific neuronal nuclei within the auditory pathway ^5–7^. We can differentiate between four to five waves, with a temporal separation of about 0.8 – 1.0 ms each^8^. Wave I is generated by the distal part of the auditory nerve (AN). Wave II reflects the projection of the cochlear nucleus (CN); Wave III is generated by the superior olivary complex (SOC), wave IV by the lateral lemniscus and inferior colliculus (IC), and lastly wave V reflecting signal transmission from the thalamus to the auditory cortex (AC) ^9–11^.

The neurotransmitter systems in the auditory brainstem circuitry are mainly glutamatergic, GABAergic, glycinergic and cholinergic ^12–14^. The CN sends glutamatergic projections to the lateral superior olive (LSO) via the medial superior olive (MSO) and medial nucleus of the trapezoid body (MNTB). MNTB neurons make glycinergic inhibitory synapses with the LSO neurons. SOC neurons send glutamatergic projections to the lateral lemniscus and the IC, targeting the medial geniculate body (MGB) in the thalamus, which sends glutamatergic projections again to the primary auditory cortex (AC)^12,13^. The descending auditory projections start from the AC and terminate in subcortical auditory centers, such as the MGB^15^ and the ventral nucleus of the trapezoid body in the auditory brainstem^16^.

While ABRs have been used extensively to assess auditory brainstem physiology^17^ and its abnormalities^18,19^, the capacity of ABRs to be modulated by pharmacological agents remains poorly understood. Therefore, we set out to test the effects of a wide range of pharmacological modulators on rodent ABRs. Here we used acute pharmacological treatments prior to the ABRs measurements. We tested the effects of enhancing the GABAergic neurotransmission in the auditory brainstem via injecting diazepam (a γ2-containing GABAA receptor enhancer), gaboxadol (a GABAA receptor agonist) or baclofen (a GABAB receptor agonist). Both GABAA and GABAB receptors are widely expressed along the different nuclei in the auditory brainstem^20–24^. A previous study emphasized the role of baclofen and diazepam as potent modulators of both the excitability of neurons in the ascending auditory pathway and the processing of auditory information by IC neurons^25^. Moreover, we used bitopertin (a non-competitive selective inhibitor of glycine transporter 1 (GlyT-1)^26^, in order to investigate the role of increased glycinergic neurotransmission on ABRs. GlyT-1 is one of the two Glycine transporters family, which work as an endogenous regulator of Glycine, but also play a key role in maintaining glycine neurotransmission homeostasis and modulating glycine levels at N-methyl-D-aspartic acid (NMDA) sites^27^. GlyT-1 are widely expressed in neuronal and glial cells^28^, among the different brain regions including the auditory brainstem^29^. We used also retigabine (a broadKv7 enhancer), which is well known to increase neuronal hyperpolarization^30^ and thus may reduce synaptic outputs in the auditory brainstem by acting on the Kv7.4 channels of the outer hair cells in the inner ear^31^. In addition, we used nicotine (a nAChR agonist) in order to inhibit excitatory output of the inner hair cells in the cochlea^32^.

Initially, we tested these compounds under the application of isoflurane, the most widely used anesthesia method for rodent ABRs^1^. In a second step, we also tested a subset of compounds under medetomidine anesthesia that may better preserve the dynamics of neural circuits and therefore could reveal compound effects different to those under isoflurane. Beside the application of different anesthesia methods, we used both Nrxn1α KO and wildtype Sprague Dawley rats in our study. Our choice was informed by previous results, showing that auditory processing is substantially impaired in Nrxn1α KO rats, and that cortical auditory responses are impacted differently by GABAergic modulation compared to their wildtype littermates^33^. Therefore, the inclusion of Nrxn1α KO Sprague Dawley rats in our current study allowed us to test if some aspects of our previous findings could be explained by functional alterations of auditory brainstem circuits.

## 2 Materials and methods

### 2.1 Animals

Experiments were conducted on adult Nrxn1α KO rats and wildtype littermates (strain: Sprague Dawley (SD)-Nrxn1<tm1sage> bred by Charles River, France. Only male rats were used. Rats were housed in groups of two, in a temperature-controlled room on a 12 h light/dark cycle with ad libitum food and water. Overall, four animal cohorts have been used, as table 1 shows.

**Table 1.** **shows the different used animal cohorts according to each pharmacological treatment and anesthesia type in both wildtype (WT) and Nrxn1α KO Sprague Dawley rats (KO)**.

| Anesthesia type | Pharmacology | Animal Cohort |
| --- | --- | --- |
| Ketamine/xylazine | No treatment -<br>Only for the Genotype comparison | Group A, WT (N=9), KO (N=9), 25 weeks old |
| Medetomidine | No treatment -<br>Only for the Genotype comparison | Group B, WT (N=12), KO (N=12), 27 weeks old |
| Isoflurane | No treatment -<br>Only for the Genotype comparison | Group B, WT (N=12), KO (N=12), 27 weeks old |
| Isoflurane | Diazepam, gaboxadol | Group C, WT (N=14), KO (N=14), 20 weeks old |
| Isoflurane | Gaboxadol | Group D, WT (N=18), KO (N=16), 20 weeks old |
| Isoflurane | Retigabine | Group D, WT (N=18), KO (N=16), 20 weeks old |
| Isoflurane | Nicotine | Group A, WT (N=14), KO (N=14), 20 weeks old |
| Isoflurane | Bitopertin | Group A, WT (N=14), KO (N=11), 20 weeks old |
| Isoflurane | Baclofen | Group A, WT (N=14), KO (N=13), 20 weeks old |
| Medetomidine | Diazepam, bitopertin, retigabine | Group B, WT (N=12), KO (N=12), 25 weeks old |

All procedures were approved by the Federal Food Safety and Veterinary Office of Switzerland and conducted in adherence to the Swiss federal ordinance on animal protection and welfare, as well as according to the rules of the Association for Assessment and Accreditation of Laboratory Animal Care International.

### 2.2 Anesthesia

isoflurane-based anesthesia started with inducing unconsciousness via isoflurane inhalation (Isofluran Baxter, Cat. no.: hdg9623, Baxter, GER), in a chamber filled with 5% isoflurane for 3 min and maintained throughout the ABRs recording at 2.5% isoflurane in medical air.

For medotidmine-based anesthesia, animals were first anesthetized via isoflurane inhalation (4% isoflurane for 4 min), and then injected with a bolus of medetomidine (0.1 mg/kg, s.c., Dorbene, Graeub, CH), followed by 1 min isoflurane inhalation at 4% to maintain anesthesia until the effect of medetomidine fully unfolded. Before starting the ABRs measurements, isoflurane inhalation was stopped for 5 min to ensure isoflurane washout. At the end of the recording, Atipamezoli (0.1 mg/kg, s.c., Alzane, Graeub, CH) was injected to reverse the sedative and analgesic effects of medetomidine. ketamine-based anesthesia was performed by i.p injection of a ketamine/xylazine mixture (80 mg/kg ketamine mixed with 5 mg/kg xylazine, i.p., Ketasol 100 with Xylasol, Graeub, CH). ABRs measurements were started 10 min after injection.

### 2.3 Pharmacology

Doses and pre-treatment times were chosen according to previously established pharmacokinetic/pharmacodynamics profiles^33–37^. Dosing was randomized using a Latin-based square design; each animal receiving every compound (or vehicle) to allow within-subject comparison. No blinding was performed. The duration of the washout phase between dosing was at least 48 hours. The control condition was represented by the administration of an equal volume of the vehicle solution (0.9% saline + 0.3% Tween20: Cat. no.: 11332465001, Sigma-Aldrich, GER). Animals were injected with diazepam (3 mg/kg, Roche Pharmaceuticals, CH), gaboxadol (10 mg/kg, Cat. no.: T101, Sigma-Aldrich, GER), retigabine (3 mg/kg, Roche Pharmaceuticals, CH), nicotine (5 mg/kg, (−) nicotine hydrogen tartrate salt, Cat. no.: SML1236, Sigma– Aldrich, GER), baclofen (5 mg/kg, Cat. no.: B5399, Sigma–Aldrich, GER), bitopertin (10 mg/kg, Roche Pharmaceuticals, CH) or vehicle solution. Intraperitoneal injection was performed 15 min before starting the ABRs measurement for all compounds, except for bitopertin, which reaches maximal exposures at around 60 minutes after application.

### 2.4 Electrophysiological recording and acoustic stimulation

Prior to the ABRs measurements, sound volume calibration was performed following the RZ6 Open Field Calibration Setup (Tucker-Davis Technologies, FL), including a signal conditioner and a 1/4-inch Prepolarized Free-field microphone (model nr. 480c02, ICP® SENSOR, PCB, NY, USA). The acoustic stimuli used in the ABRs assessment consisted of 512 click tones responses with 50 ms inter-stimulus interval. Each click is a broadband mono-phasic square wave signal (0.1 ms). The clicks were presented at a rate of 21 clicks/s, at different sound levels (90, 80, 70, 60, 50, 40, 30, 20, 10 dB SPL), starting with the highest stimulus intensities, in line with established protocols ^33^. The ABRs measurements were conducted in a sound-attenuating and electrostatically grounded chamber. Body temperature of anesthetized animals (see above) was maintained at 37° C using a thermic heating pad (Kent Scientific Corporation, CN, USA). Click-tones were generated with a multi field speaker (MF1, Tucker-Davis Technologies, FL, USA) connected to a RZ6-A-1 input/output processor (Tucker-Davis Technologies, FL, USA). The speaker was positioned 10 cm from the animal’s right ear. ABRs signals were recorded with 13 mm subdermal needle electrodes (Cat. no.: NS-s83018-r9-10, Rochester, Coral Springs, FL, USA), with the signal electrodes placed on the vertex and reference and ground electrodes placed under the ipsi- and contralateral ear, respectively, connected to a RA4PA preamplifier/digitizer and RA4LI low impedance head stage (Tucker-Davis Technologies, FL, USA). Signals were acquired using the following settings: 12 kHz sampling rate, 5 kHz low pass, 100 Hz high pass, 50 Hz notch, using the BioSigRZ software (version 5.5, TDT, FL, USA).

### 2.5 Data processing and analysis

Data analysis was performed as previously described ^33^. In brief, in a pre-processing step ABRs data was normalized to its pre-stimulus baseline. Resulting ABRs waveforms were statistically tested for differences between conditions (see Statistical testing).

### 2.6 Statistical testing

Statistical testing was performed with paired or unpaired cluster-based permutation tests (CBPT) depending on the condition, using custom *Python* scripts. In brief, first CBPT performs individual t-tests (two-tailed, significance level set to p <0.05) for each data point. The resulting clusters are then tested for significance by comparing the summed t-values of the initial clusters with summed t-values of clusters obtained from permuted data (here, shuffling over the time domain) over many iterations (N = 1000 permutations, significance threshold: p <0.05), thereby correcting for multiple comparisons. We visualize both cluster types (with permutation testing: black bars above graphs; and w/o permutation: grey bars, indicating statistical trends). Given that qualitatively no apparent outliers were present, no specific test was performed for outlier detection. No exclusion criteria were predetermined, and no animals were excluded from the statistical analysis. For one animal under one condition in the pharmacology study (nicotine, 5.0 mg/kg), missing vehicle data was input by averaging the respective data points of all other animals under this condition, to allow for paired analyses. In all plots, data is displayed as mean ± standard error of mean (shaded area), except mentioned otherwise.

## 3 Results

### 3.1 Auditory brainstem responses are similar for adult Nrxn1α KO Sprague Dawley rats and wildtype littermates under different anesthetics

First, we asked whether Nrxn1α KO Sprague Dawley rats show alterations in their ABRs compared to wildtype littermates. To mitigate the risk that putative genotypic differences are missed due to the effects of a particular anesthesia, we chose to perform ABRs recordings under three different types of anesthesia. We found that under all conditions, ABRs of Nrxn1α KO animals largely resembled those of their wildtype littermates (Fig. 1 and Supplementary Fig. 1 and 2). Except for statistically significant differences in the very late components of the ABRs elicited at 80 dB under medetomidine (Supplementary Fig. 2B; time window 5.4 – 6.5 ms, d = -1.23, p = 0.028 and time window 7 – 8.5 ms, d = 1.12, p = 0.012). While we did not notice remarkable genotypic differences, we noted that ABRs were affected by choice of anesthetic. In fact, the overall amplitude of the deflections appeared strongest under ketamine/xylazine, and ABRs under both ketamine/xylazine and medetomidine showed better-resolved individual waves compared to isoflurane.

**Figure 1.**
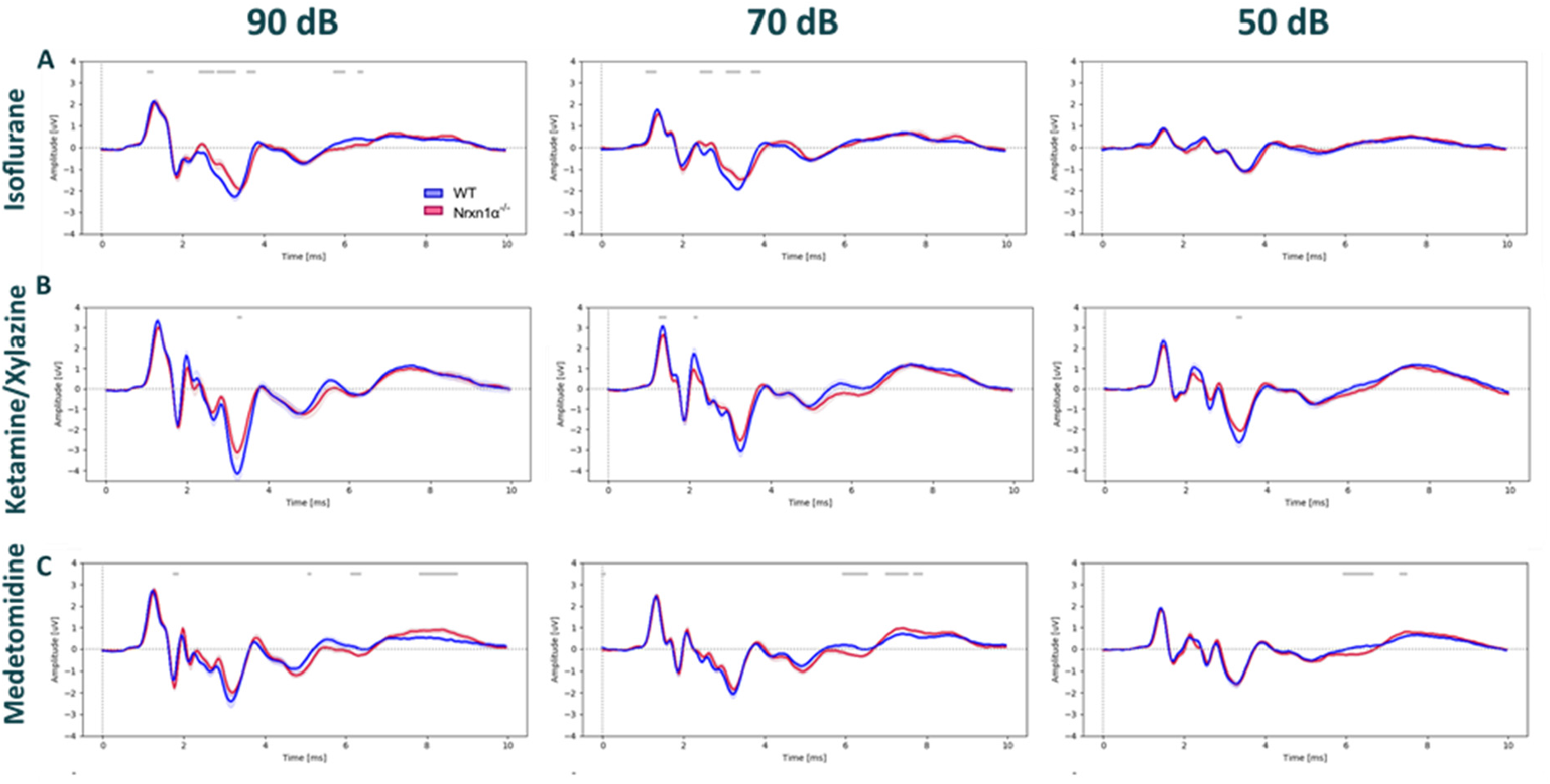
Comparison of auditory brainstem responses between Nrxn1α KO Sprague Dawley and wild-type littermates rats. ABRs waveforms across different stimulus intensities (90, 70, 50 dB) under A) isoflurane, B) ketamine/xylazine and C) medetomidine anesthesia. Recordings from the WT are in blue (N=12) and Nrxn1α KO in red (N=12). Data displayed as mean ± SEM, was tested with unpaired CBPT. No robust significant differences were found between genotypes across anesthesia methods. Grey bars above the graphs indicate clusters of significant differences before CBPT-based correction for multiple comparisons, i.e. indicating statistical trends.

### 3.1 ABRs are largely resistant to pharmacological modulators under isoflurane anesthesia

Next, we assessed how pharmacological agents that modulate distinct neurotransmitter systems impact ABRs in both wildtype (Fig. 2 and Supplementary Fig. 3) or Nrxn1α KO Sprague Dawley rats (Fig. 3 and Supplementary Fig. 4). In our first set of experiments, we used isoflurane anesthesia, as it is arguably the most-widely used choice for rodent ABRs measurements. In order to investigate the effects of increasing GABAergic neurotransmission, we tested diazepam at 3 mg/kg (a γ2-containing GABA_A_ receptor enhancer; Fig. 2A and 3A), gaboxadol at 10 mg/kg (α4/6δ-containing GABA_A_ receptor agonist; Fig. 2B and 3B) and baclofen at 5 mg/kg (a GABA_B_ receptor agonist; Fig. 2C and 3C). To augment glycinergic neurotransmission we used bitopertin at 10 mg/kg (a glycine transporter 1 inhibitor; Fig. 2D and 3D). We used retigabine at 3 mg/kg (a pan-K_v_7 enhancer; Fig. 2E and 3E) to increase neuronal hyperpolarization and, therefore, to overall reduce synaptic outputs. nicotine was used at 5 mg/kg (a nAChR agonist; Fig. 2F and 3F) in order to inhibit output of inner hair cells of the cochlea. Interestingly, we found that, compared to the vehicle control, none of the applied pharmacological agents clearly impacted ABRs in either wildtype or Nrxn1α KO Sprague Dawley rats. The only statistically significant effects were observed with nicotine on ABRs elicited at 90 dB and with retigabine on ABRs elicited at 80 dB. nicotine showed a modulation of the very late components of the ABRs in both wildtype (Fig. 2F; time window 5.4 – 6.25 ms time window, d = -1.27, p = 0.037) and Nrxn1α KO Sprague Dawley rats (Fig. 3F; time window 6.6 – 7.9 ms; d = 0.97, p = 0.009), while retigabine only affected ABRs of Nrxn1α KO Sprague Dawley rats (Supplementary Fig. 4C, time window 3.6 – 5.9 ms, d = - 0.84, p = 0.01).

**Figure 2.**
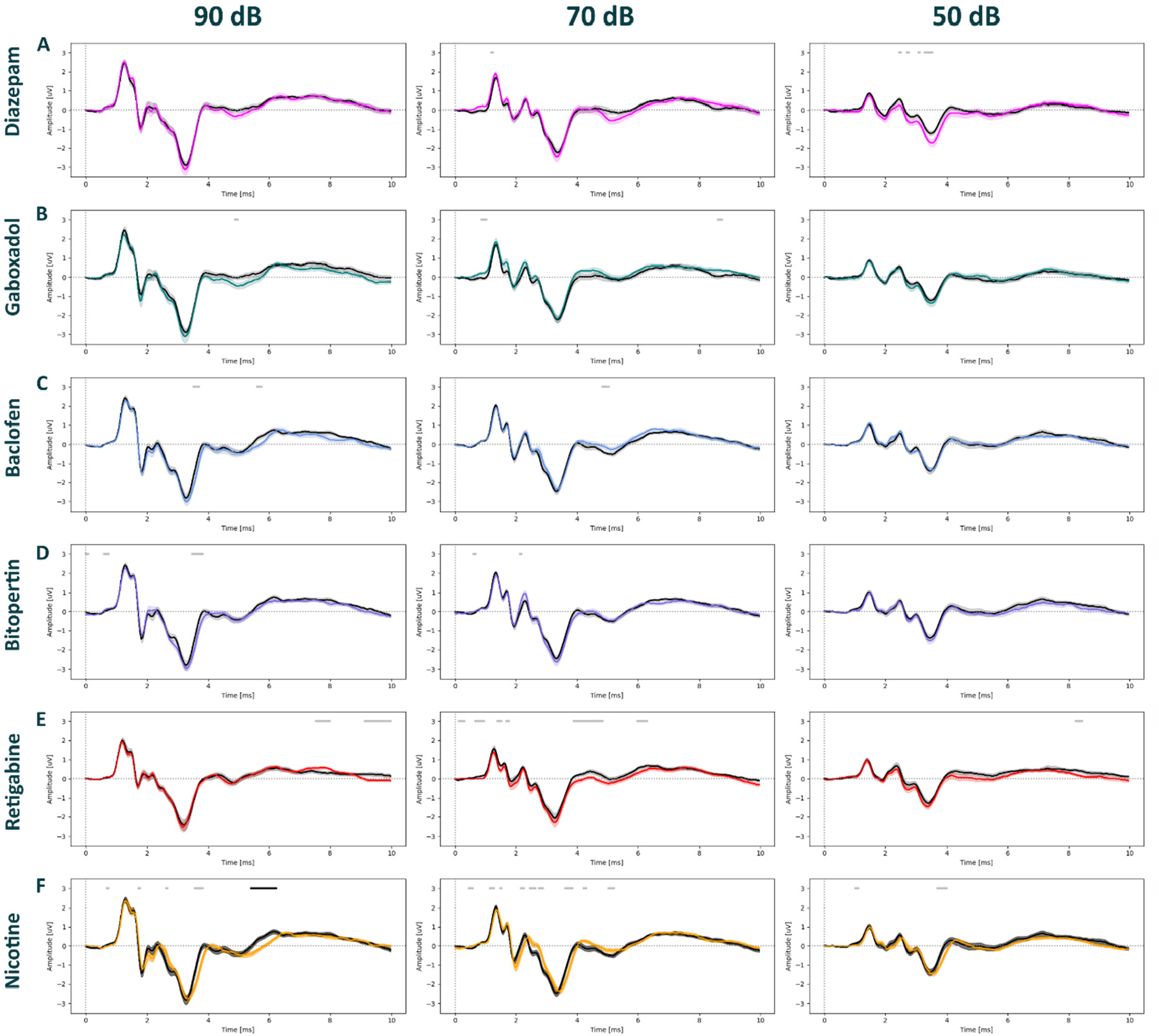
Auditory brainstem responses post pharmacological treatment in WT Sprague Dawley rats under isoflurane anesthesia. ABRs waveforms across different stimulus intensities (90, 70, 50 dB) post intraperitoneal injection with diazepam (3 mg/kg) in magenta; (N=14), gaboxadol (10 mg/kg) in teal; (N=14), baclofen (5 mg/kg) in blue; (N=14), bitopertin (10 mg/kg) in purple, retigabine (3 mg/kg) in red; (N=18), nicotine (5 mg/kg) in yellow; (N=14),; (N=14), or vehicle solution in black (0.9% saline + 0.3% Tween). Within each experimental block, dosing was counterbalanced, and applied 15 min prior to the ABRs recording for all compounds, except for bitopertin (60 min pre-treatment time). The Black bars above the graphs indicate clusters of significant differences between conditions. The Gray bars indicate clusters that have not reached significance threshold post-permutations. Data displayed as mean ± SEM.

**Figure 3.**
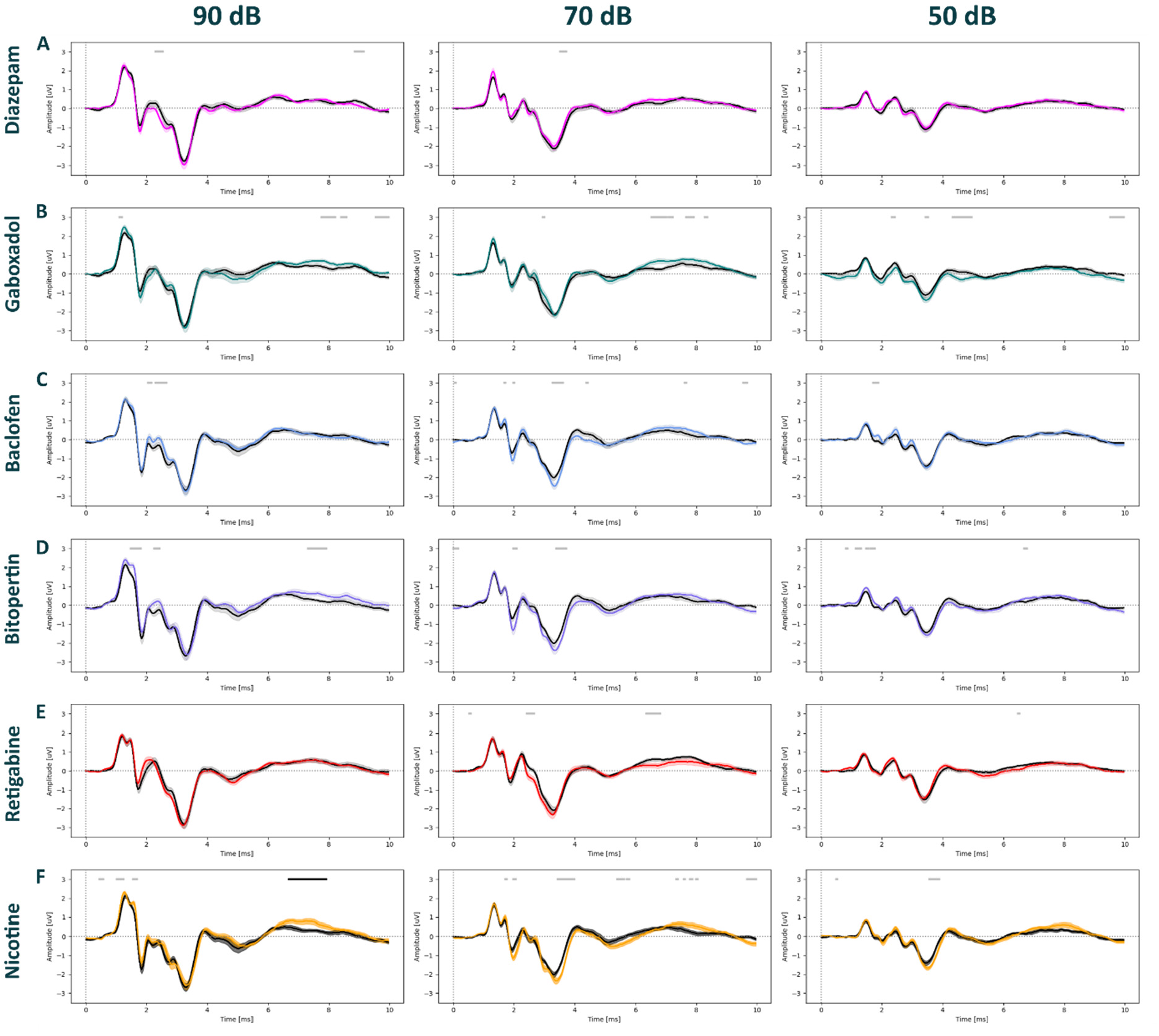
Auditory brainstem responses post pharmacological treatments in Nrxn1α Sprague Dawley rats under isoflurane anesthesia. ABRs waveforms across different stimulus intensities (90, 70, 50 dB) post intraperitoneal injection with diazepam (3 mg/kg) in magenta; (N=14), gaboxadol (10 mg/kg) in teal; (N=14), baclofen (5 mg/kg) in blue; (N=13), bitopertin (10 mg/kg) in purple; (N=11), retigabine (3 mg/kg) in red; (N=16), nicotine (5 mg/kg) in yellow; (N=14), or vehicle solution in black (0.9% saline + 0.3% Tween). Within each experimental block, dosing was counterbalanced, and applied 15 min prior to the ABRs recording for all compounds, except for in bitopertin (60 min pre-treatment time). The Black bars above the graphs indicated CBPT clusters of significant differences within subjects, i.e., between conditions. The Gray bars indicate clusters that have not reached significance threshold post-permutations. Data displayed as mean ± SEM.

### 3.3 ABRs are largely resistant to pharmacological modulations under medetomidine anesthesia

With the lack of pharmacological modulation observed under isoflurane, we next tested if ABRs could be modulated more clearly under medetomidine, a widely-used anesthetic in functional imaging that is discussed to preserve better network dynamics as compared to isoflurane or ketamine. To test this hypothesis, we focused on the three compounds diazepam (Fig. 4A, 5A), bitopertin (Fig. 4B, 5B) and retigabine (Fig. 4C, 5C). Similar to our observations under isoflurane, pharmacological modulation did not alter ABRs of both wildtype (Fig. 4 and Supplementary Fig. 5) and Nrxn1α KO Sprague Dawley rats (Fig. 5 and Supplementary Fig. 6) under medetomidine. The only statistically significant difference was found for retigabine in wildtype animals, reducing the amplitude of late components of ABRs elicited at 40 dB (Supplementary fig. 5C; time window 3.8 – 5.5 ms, d = -1.44, p = 0.012; and time window 5.6 – 7 ms, d = -1.68, p = 0.013).

**Figure 4.**
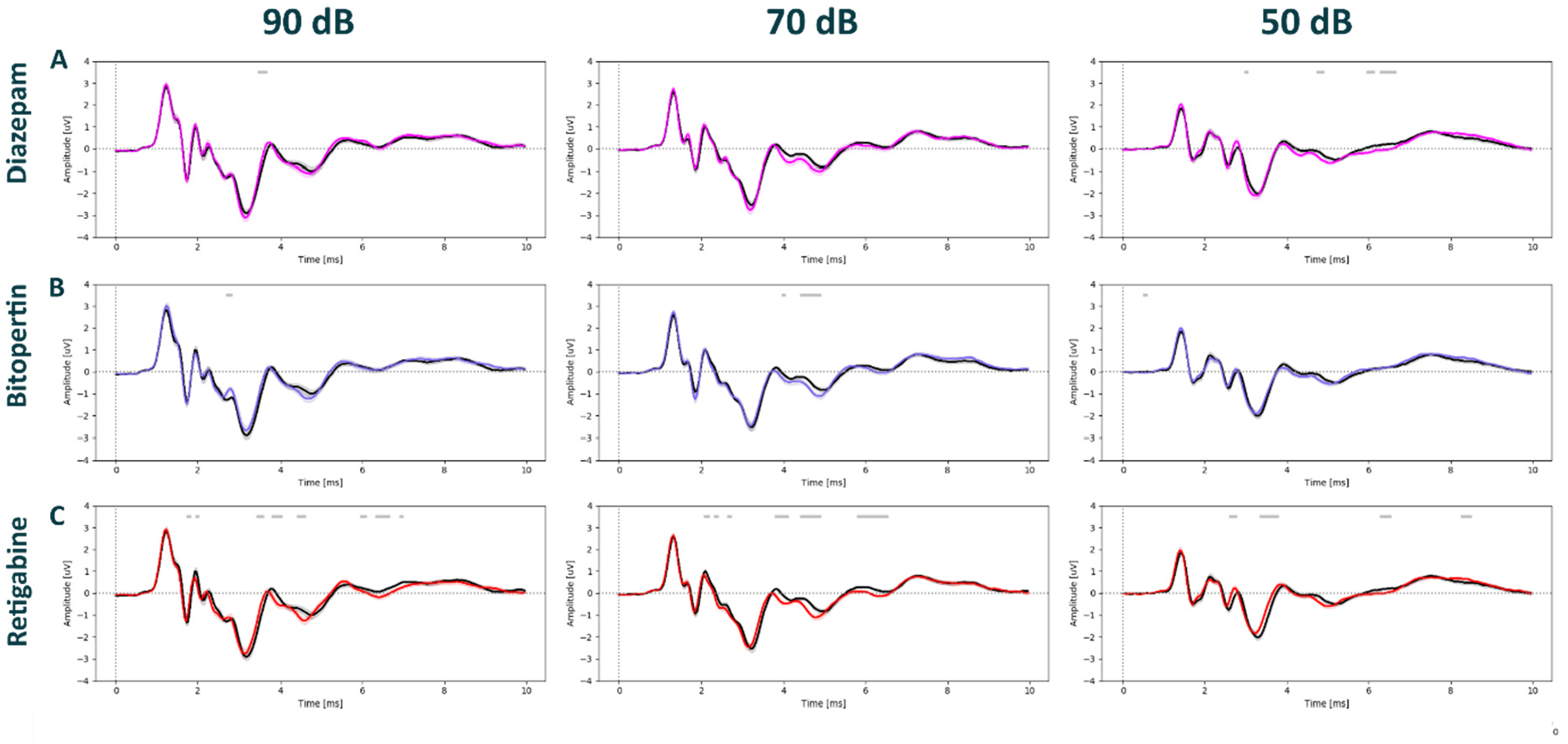
Auditory brainstem responses post pharmacological treatment in WT Sprague Dawley rats under medetomidine anesthesia. ABRs waveforms across different stimulus intensities (90, 70, 50 dB) post intraperitoneal injection with diazepam (3 mg/kg) in magenta; (N=12), bitopertin (10 mg/kg) in purple; (N=12), retigabine (3 mg/kg) in red; (N=12), or vehicle solution in black (0.9% saline + 0.3% Tween). Within each experimental block, dosing was counterbalanced, and applied 15 min prior to the ABRs recording for all compounds, except for in bitopertin (60 min pre-treatment time). Data displayed as mean ± SEM, was tested with unpaired CBPT. No robust significant differences were found between genotypes across anesthesia methods. Grey bars above the graphs indicate clusters of significant differences before CBPT-based correction for multiple comparisons, i.e., indicating statistical trends.

**Figure 5.**
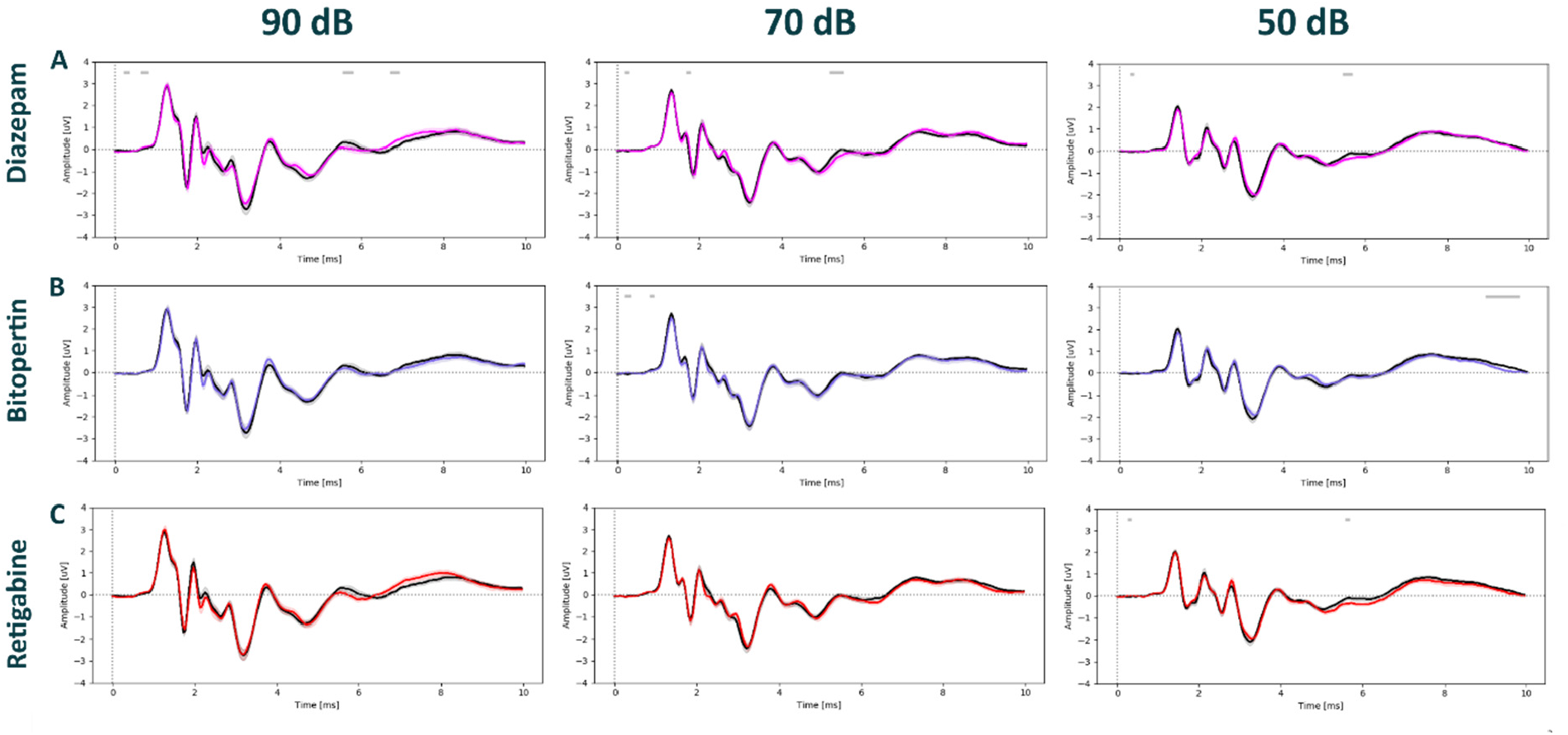
Auditory brainstem responses post pharmacological treatment in Nrxn1α KO Sprague Dawley rats under medetomidine anesthesia. ABRs waveforms across different stimulus intensities (90, 70, 50 dB) post intraperitoneal injection with diazepam (3 mg/kg) in magenta; (N=12), bitopertin (10 mg/kg) in purple; (N=12), retigabine (3 mg/kg) in red; (N=12), or vehicle solution in black (0.9% saline + 0.3% Tween). Within each experimental block, dosing was counterbalanced, and applied 15 min prior to the ABRs recording for all compounds, except for in bitopertin (60 min pre-treatment time). Data displayed as mean ± SEM, was tested with unpaired CBPT. No robust significant differences were found between genotypes across anesthesia methods. Grey bars above the graphs indicate clusters of significant differences before CBPT-based correction for multiple comparisons, i.e., indicating statistical trends.

## 4 Discussion

The current study explored the impact of different anesthetics and pharmacological tool compounds in wildtype and Nrxn1α KO Sprague Dawley rats and shows for the first time that rat ABRs are unaffected by diverse pharmacological modulators.

First, by using the three most widely-used anesthetics for rodents, we confirmed that ABRs without additional pharmacological intervention are similar between adult Nrxn1α KO Sprague Dawley rats and their wildtype littermates. Our results are in line with our previous studies that probed ABRs in adult wildtype and Nrxn1α KO Sprague Dawley rats under isoflurane anesthesia only^33^. Our current study expands this finding by demonstrating the lack of genotypic differences also under ketamine/xylazine and medetomidine anesthesia. This finding is important, since previous studies showed that the choice of anesthesia (e.g., isoflurane vs. ketamine/xylazine) significantly affected ABRs characteristics^38^, raising the possibility that genotypic differences may be missed with just using one type of anesthesia with a specific mode of action. isoflurane and ketamine/xylazine (the two most widely-used anesthetics for rodents ABRs^1^) share many molecular targets, including glycine receptors^39^, GABAA^39–42^ and GABAB receptors^43,44^, glutamate receptors^45–47^ (including NMDA receptors^48–50^), and nAChR receptors^51,52^. All these receptors are widely expressed in the brainstem and along the auditory pathway^12,13^. Any changes in these neurotransmitter systems may affect the transmission of auditory information from the cochlear to higher brain areas^51^. Indeed, Santarelli et al. showed that the latencies of ABRs waves are significantly increased during isoflurane anesthesia, in comparison to awake ABRs in Sprague Dawley rats^53^. These differences could be due to isoflurane reducing the glutamatergic neurotransmission at pre- and postsynaptic sites of inner hair cells^53^ or by augmenting GABAergic inhibition within the auditory brainstem circuits. While similar circuit engagement can be expected with ketamine/xylazine, Ruebhausen et al. showed that isoflurane elevates hearing thresholds by around 30 dB more than ketamine/xylazine-based anesthesia^38^. This could be due to an additional effect of isoflurane by increasing blood flow to the brainstem^54^ and decreasing tissue perfusion and glutamate release^55^, which potentially reduces stimulus-driven activity^38^. As an alternative to isoflurane or ketamine/xylazine we used medetomidine, an α2-adrenoceptor agonist, that is a common choice for fMRI studies as it preserved the dynamics of the brain better than α-chloralose or isoflurane^56,57^. Indeed, previous studies show that medetomidine administration only marginally influences auditory-evoked potentials, picked up in the midbrain^58^ and in the cortex^59^. Other studies show that dexmedetomidine, a medetomidine isomer, demonstrated a minimal effect on ABRs in children^60^ and it could be a better alternative for the commonly used oral chloral hydrate sedation^61^.

A key point of the current study is that testing a diverse set of pharmacological modulators showed either none or only marginal effects on the ABRs signatures. This is surprising since the tool compounds and doses used clearly engage receptors that are involved in signal transmission within auditory brainstem circuits. In fact, only nicotine and retigabine treatment led to significant, but small effects in the ABR. The effects of nicotine were confined to the very late phase of the ABR, resembling the activation of higher-order brain regions, and only at 90 dB stimulus intensity. While the major targets of nicotine (nACh receptors) are expressed at inner hair cells to regulate their sensitivity^62^, no effects on the very early components of the ABRs were evident. Therefore, our data argue for the action of nicotine on higher-order brain circuits to alter auditory processing^63^. For retigabine, we observed slightly reduced amplitudes of late components of the ABRs at 80 dB, but not at 90 dB or at 70 dB. The volume-specific effect challenges the robustness and interpretability of the finding. More importantly, the fact that retigabine enhances voltage-gated potassium channels (such as Kv7.4) expressed in the auditory brainstem^64^, but does not clearly affect the ABR, highlights yet again the resistance of ABRs to pharmacological modulation. Our findings are in line with previous studies, showing a lack of ABRs and hearing threshold modulation with retigabine^65^. Beyond our own findings with other compounds (such as benzodiazepine, baclofen or bitopertin), the notion of a more general issue with pharmacological modulation of ABRs, is further supported by other rodent studies, demonstrating the lack of ABRs modulation with opioids^66^. This is different to earlier studies demonstrating that theophylline^67^ or cocaine^68^ change ABRs characteristics likely due to ototoxic rather than neuromodulatory effects.

An intuitive explanation for the lack of pharmacological modulation of ABRs in rodents is the “masking” effects of anesthesia, which may either block the target receptors and/or reduce neuronal dynamics to the extent that does not allow for further pharmacological modulation. We mitigated this caveat by using diverse anesthetic protocols, including medetomidine which largely preserves network dynamics. Further support for the resistance of ABRs to be modulated pharmacologically comes from human and non-human primate studies which allow awake ABRs experiments. In this context, Samra et al. showed in awake rhesus monkeys that neither Scopolamine nor Morphine intravenous injection could modulate the ABRs waves^69^. In addition, studies in humans report no effects of anesthetics agents, and drugs such as Benzodiazepines, Propofol, and ketamine on ABRs^2,70^.

Independent of these considerations, our study suggests that rodent ABRs measurements are unsuited to testing auditory circuit modulation by diverse pharmacology. This conclusion is critical for drug development programs that aim to tackle auditory processing deficits, such as in psychiatric and neurodevelopmental disorders, where sensory abnormalities might stem from early-life disruption of auditory brainstem circuits^3^.

## Supporting information

Marashli et al., 2023 - Supplementary Material

## 6 Acknowledgment

We would like to acknowledge Marie Bainier for the excellent technical assistance. Furthermore, we would like to thank the Roche Innovation Centre M.Sc. Internship Program for the funding of Samuel Marashli.

## 7 Conflicts of interest

PJ and RLR were under employment by the company F. Hoffmann-La Roche (Roche). The funder provided support in the form of salaries for authors but did not have any additional role in the study design, data collection, analysis, decision to publish, or manuscript preparation. This does not alter the authors’ adherence to all the journal policies on sharing data and materials.

## 8 Data availability

The datasets generated during and/or analysed during the current study are available from the corresponding author on reasonable request.

