## Supplementary material for "Auditory brainstem responses are resistant to pharmacological modulation in Sprague Dawley wildtype and Neurexin1α knockout rats": Marashli et al., 2023 - Supplementary Material

### Supplementary information

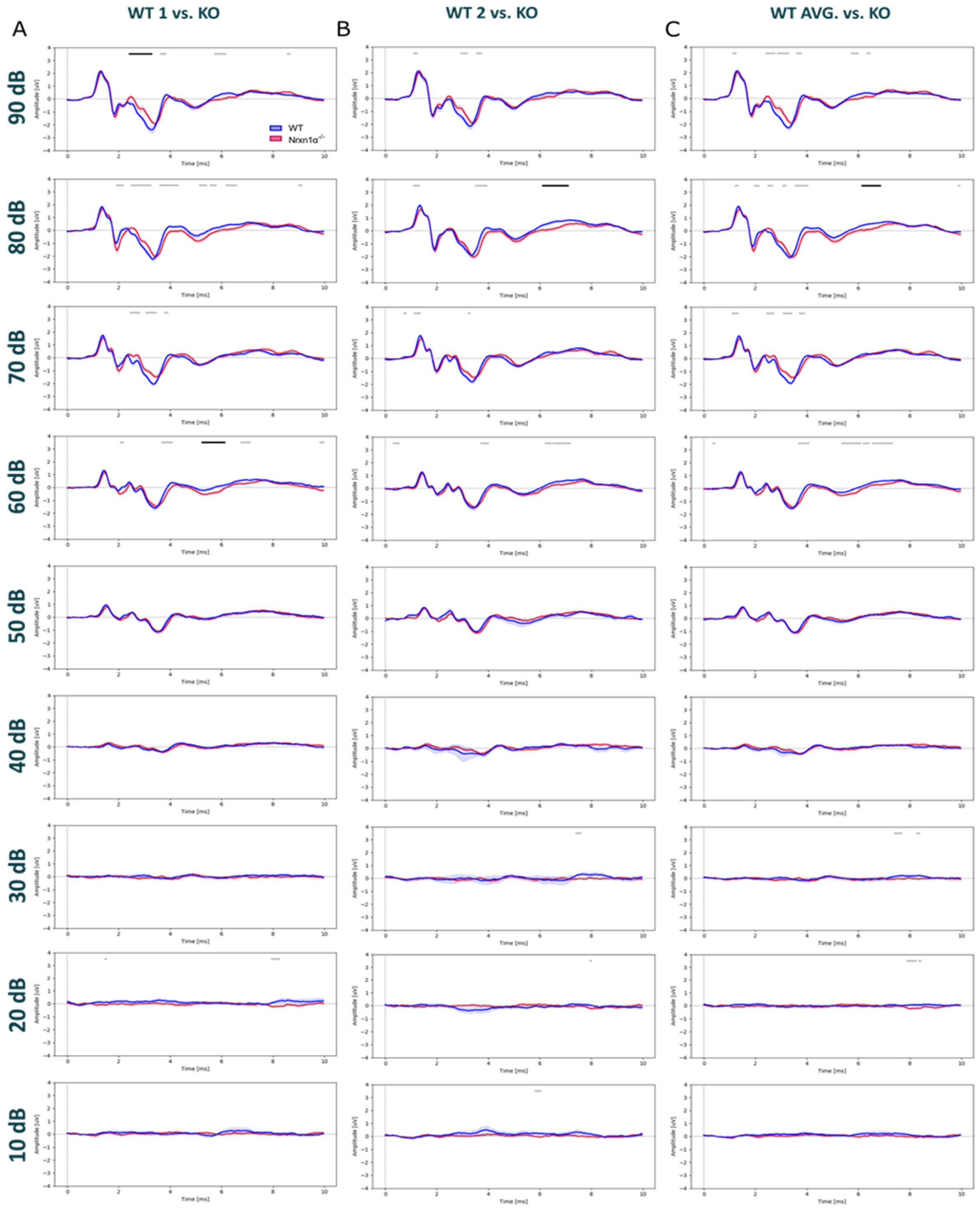


**Supplementary Figure 1: Comparison of auditory brainstem responses between Nrxn1α KO and wild-type littermates Sprague Dawley rats under isoflurane**

ABRs waveforms across different stimulus intensities (90, 80, 70, 60, 50, 40, 30, 20, 10 dB) under isoflurane-based anesthesia. Recordings from the WT are in blue (N=12) and Nrxn1α KO in red (N=12) at 27 weeks old. A) First recordings of WT in comparison with KO recordings, B) Second recordings of WT in comparison with KO recordings, C) Averaged recordings of each animal in comparison with KO recordings. Unpaired CBPT revealed significant differences between clusters following 1000 permutations ($p<0.05$) and displayed in Black bars above the graphs. The Gray bars indicate clusters that have not reached significance threshold post-permutations. Data displayed as mean ± SEM.

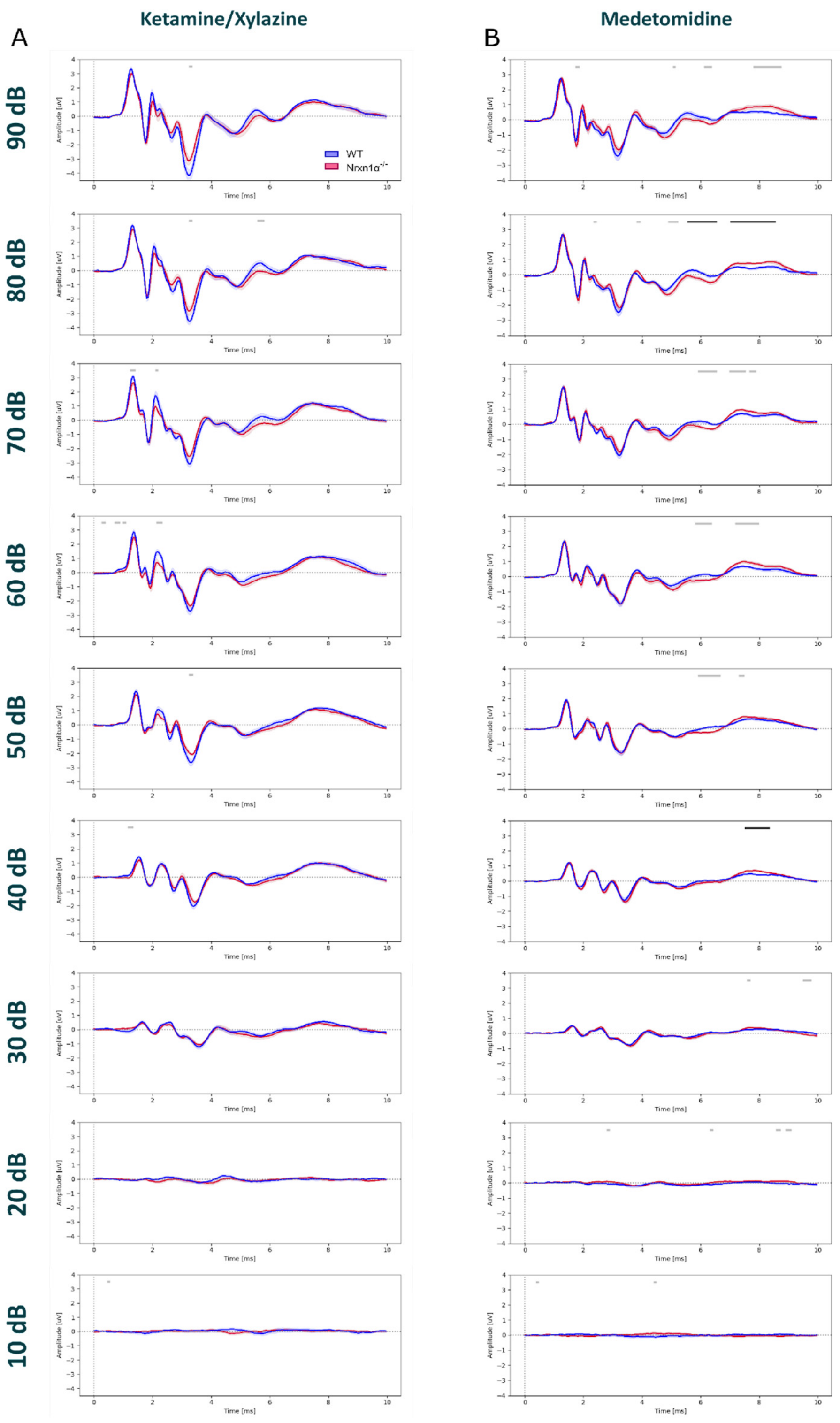


**Supplementary Figure 2: Comparison of auditory brainstem responses between Nrxn1α KO and wild-type littermates Sprague Dawley rats under ketamine/xylazine or medetomidine anesthesia**

ABRs waveforms across different stimulus intensities (90, 80, 70, 60, 50, 40, 30, 20, 10 dB). Recordings from the WT are in blue and Nrxn1α KO Sprague Dawley rats in red. A) ABRs recordings of WT (N=9) in comparison with KO (N=9) at week 25 under ketamine/xylazine based anesthesia B) ABRs recordings of WT (N=12) in comparison with KO (N=12) at week 27 under medetomidine based anesthesia. Unpaired CBPT revealed significant differences between clusters following 1000 permutations ($p<0.05$) and displayed in Black bars above the graphs. The Gray bars indicate clusters that have not reached significance threshold post-permutations. Data displayed as mean ± SEM.

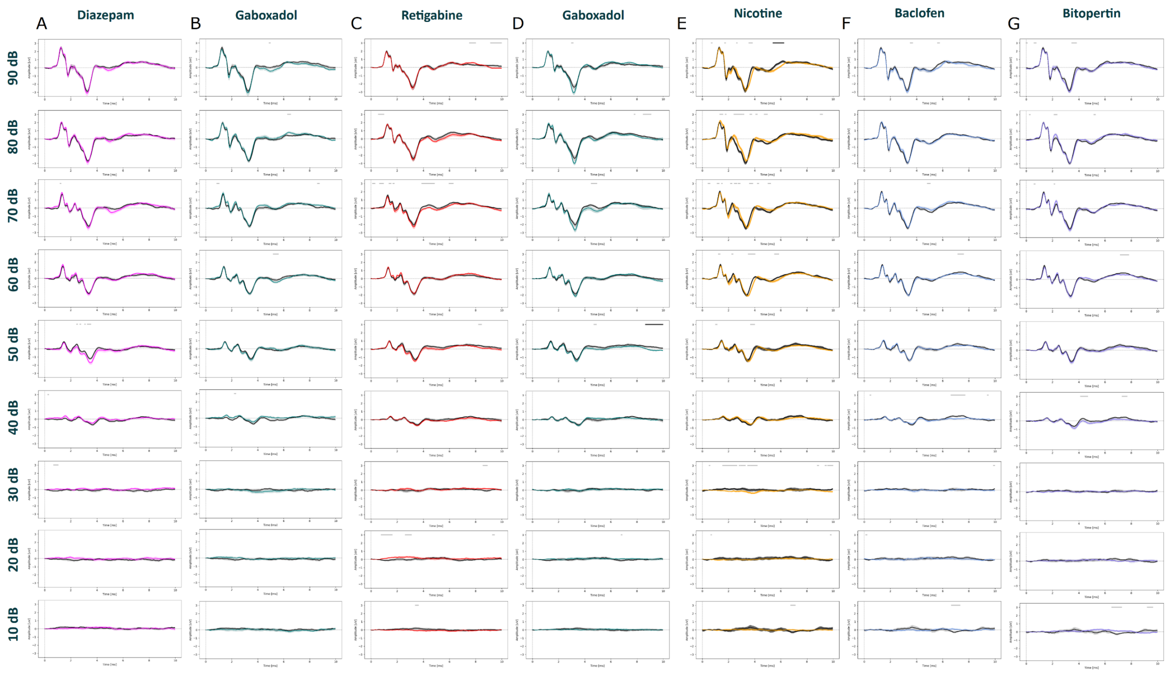


**Supplementary Figure 3: Comparison of auditory brainstem responses between pharmacological modulations and vehicle in wild-type Sprague Dawley rats under isoflurane anesthesia**

ABRs waveforms across different stimulus intensities (90, 80, 70, 60, 50, 40, 30, 20, 10 dB) post i.p. injections with different pharmacological compounds under isoflurane in WT Sprague Dawley rats. Animals (20 weeks old) were injected with; diazepam in magenta (3 mg/kg, N=14), gaboxadol in teal (10 mg/kg, (N=14)), retigabine in red (3 mg/kg, N=18)), nicotine in yellow (5 mg/kg, N=14)), baclofen in blue (5 mg/kg, N=14), bitopertin in purple (10 mg/kg, (N=14), or vehicle solution in black (0.9% saline + 0.3% Tween). Within each experimental block, dosing was counterbalanced, with the order of animals staying the same, and applied 15 min prior to the ABRs recording for all compounds, except in bitopertin (60 min). Paired CBPT revealed significant differences between clusters following 1000 permutations ($p<0.05$) and displayed in Black bars above the graphs. The Gray bars indicate clusters that have not reached significance threshold post-permutations. Data displayed as mean ± SEM. D) repeated gaboxadol treatment in different animal cohorts (the same animal cohort of retigabine) (3 mg/kg, (N=18), not shown as main figure.

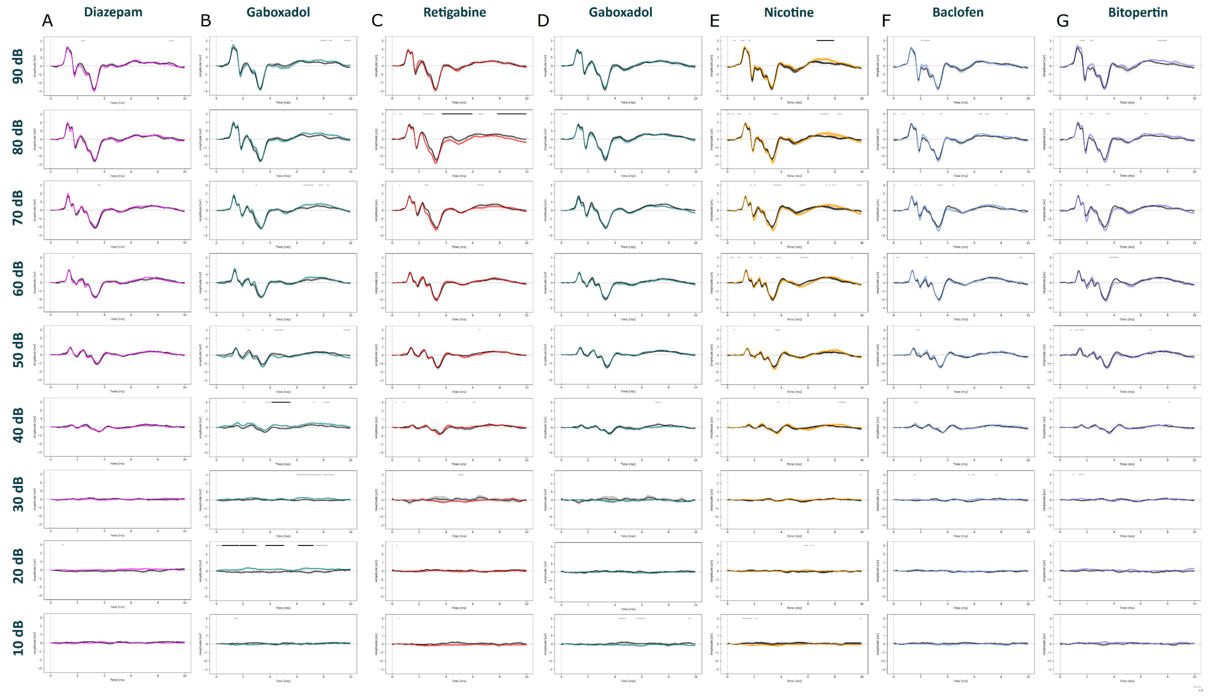


**Supplementary Figure 4: Comparison of auditory brainstem responses between pharmacological modulations and vehicle in Nrxn1α KO Sprague Dawley rats under isoflurane anesthesia**

ABRs waveforms across different stimulus intensities (90, 80, 70, 60, 50, 40, 30, 20, 10 dB) post i.p. injections with different pharmacological compounds under isoflurane in Nrxn1α KO Sprague Dawley rats. Animals (20 weeks old) were injected intraperitoneally with; diazepam in magenta (3 mg/kg, (N=14)), gaboxadol in teal (10 mg/kg, (N=14)) retigabine in red (3 mg/kg, (N=16)), nicotine in yellow (5 mg/kg, (N=14)), baclofen in blue (5 mg/kg, (N=13)), bitopertin in purple (10 mg/kg, (N=11)), or vehicle solution in black (0.9% saline + 0.3% Tween). Within each experimental block, dosing was counterbalanced, with the order of animals staying the same, and applied 15 min prior to the ABRs recording for all compounds, except in bitopertin (60 min). Paired CBPT revealed significant differences between clusters following 1000 permutations ($p<0.05$) and displayed in Black bars above the graphs. The Gray bars indicate clusters that have not reached significance threshold post-permutations. Data displayed as mean ± SEM. D) repeated gaboxadol treatment in different animal cohorts (the same animal cohort of retigabine) (3 mg/kg, (N=16), not shown as main figure.

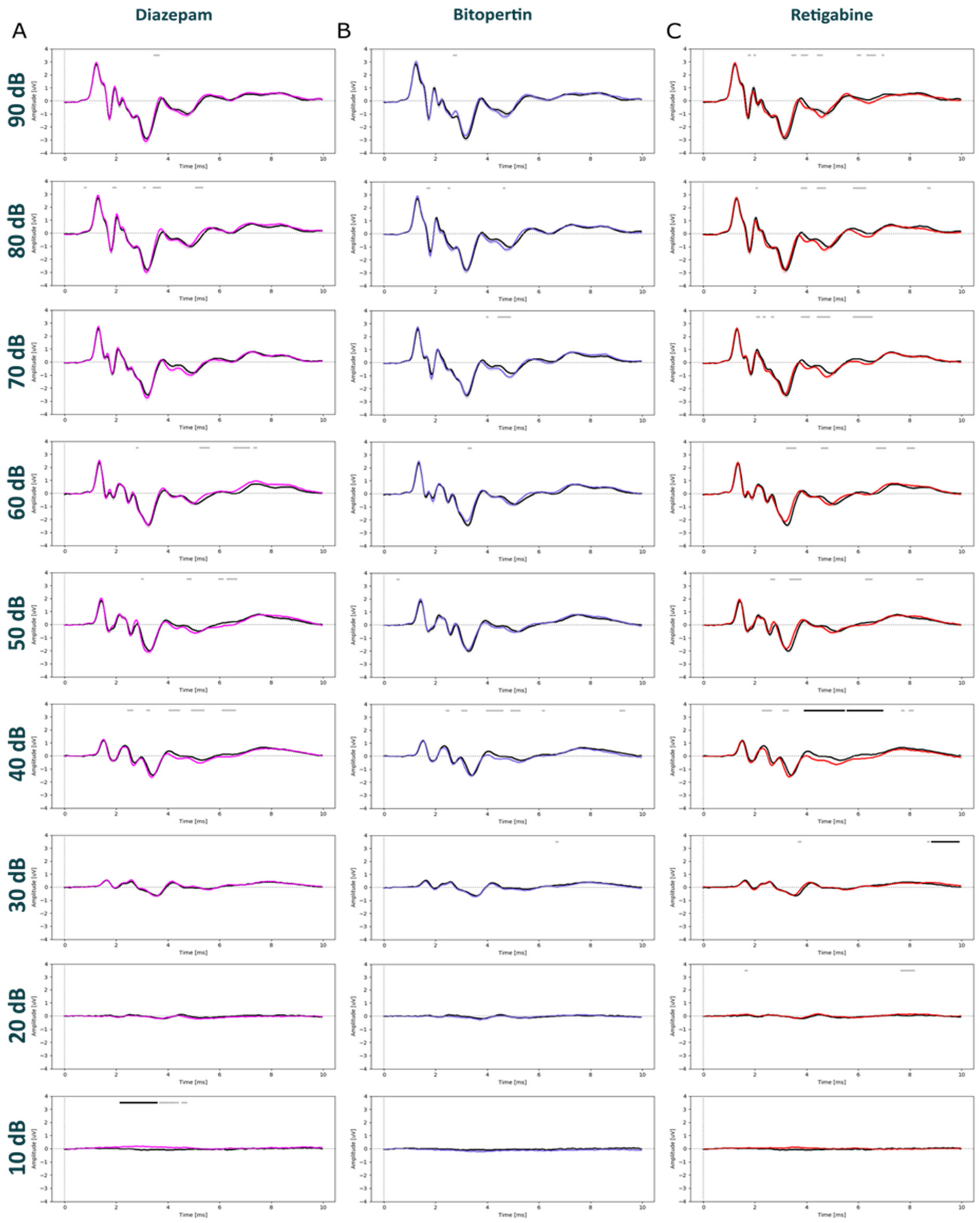


**Supplementary Figure 5: Comparison of auditory brainstem responses between pharmacological modulations and vehicle in wild-type Sprague Dawley rats under medetomidine anesthesia**

ABRs waveforms across different stimulus intensities (90, 80, 70, 60, 50, 40, 30, 20, 10 dB) post i.p. injections with different pharmacological compounds under medetomidine in WT Sprague Dawley rats. diazepam (3 mg/kg) in magenta; (N=12), bitopertin (10 mg/kg) in purple; (N=12), retigabine (3 mg/kg) in red; (N=12), or vehicle solution in black (0.9% saline + 0.3% Tween). Within each experimental block, dosing was counterbalanced, and applied 15 min prior to the ABRs recording for all compounds, except in bitopertin (60 min). The Black bars above the graphs indicated CBPT significant differences within-subjects, i.e., between the vehicle and nicotine treatments ($p<0.05$). The Gray bars indicate clusters that have not reached significance threshold post-permutations. Data displayed as mean ±.

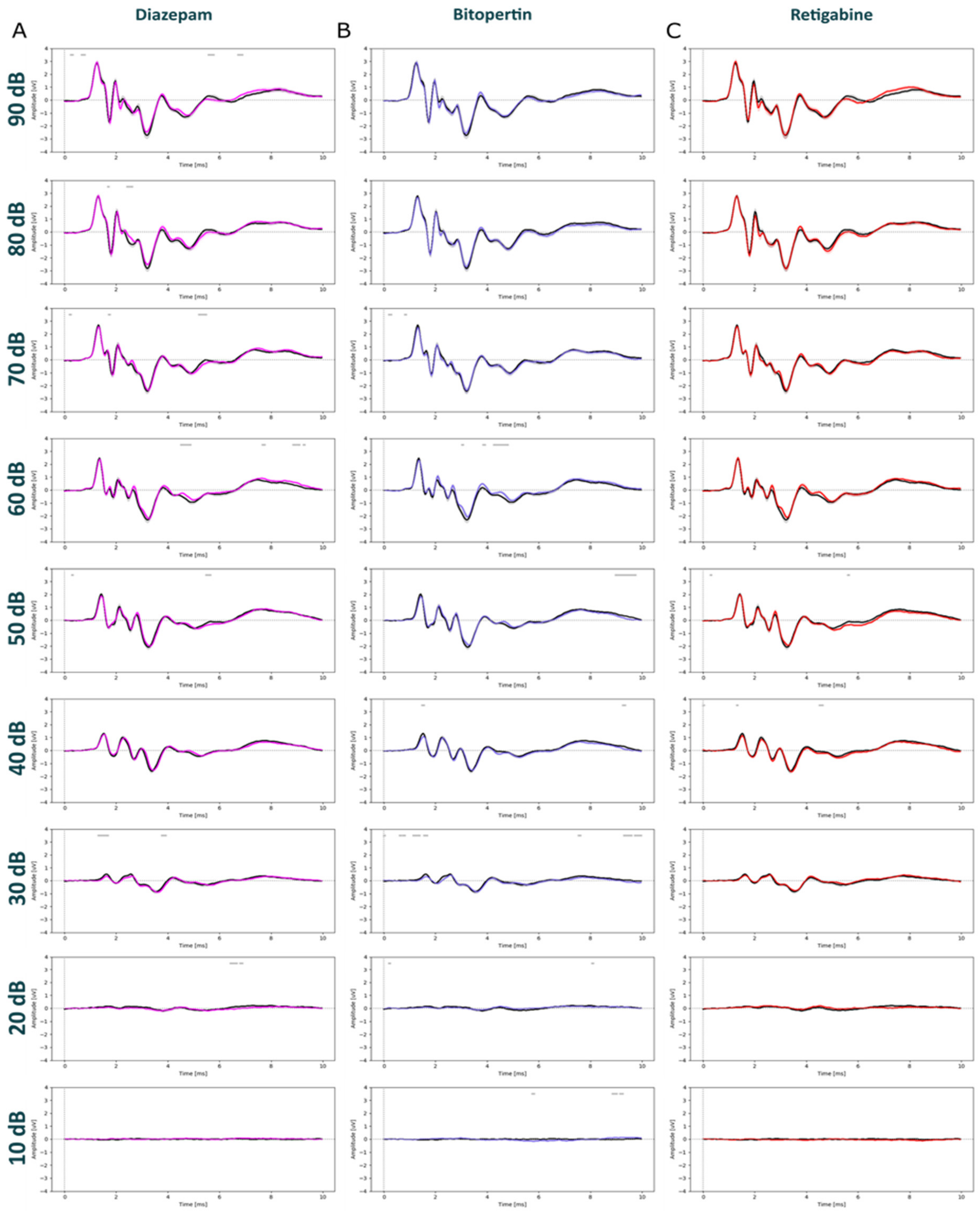


**Supplementary Figure 6: Comparison of auditory brainstem responses between pharmacological modulations and vehicle in Nrxn1α KO Sprague Dawley rats under medetomidine anesthesia**

ABRs waveforms across different stimulus intensities (90, 80, 70, 60, 50, 40, 30, 20, 10 dB) post i.p. injections with different pharmacological compounds under medetomidine in Nrxn1α KO Sprague Dawley rats. diazepam (3 mg/kg) in magenta; (N=12), bitopertin (10 mg/kg) in purple; (N=12), retigabine (3 mg/kg) in red; (N=12), or vehicle solution in black (0.9% saline + 0.3% Tween). Within each experimental block, dosing was counterbalanced, and applied 15 min prior to the ABRs recording for all compounds, except in bitopertin (60 min). The Black bars above the graphs indicated CBPT significant differences within-subjects, i.e., between the vehicle and nicotine treatments ($p<0.05$). The Gray bars indicate clusters that have not reached significance threshold post-permutations. Data displayed as mean ± SEM.
